# SARTools: a DESeq2- and edgeR-based R pipeline for comprehensive differential analysis of RNA-Seq data

**DOI:** 10.1101/021741

**Authors:** Hugo Varet, Jean-Yves Coppée, Marie-Agnès Dillies

**Affiliations:** Center for Bioinformatics, Biostatistics and Integrative Biology (C3BI), Institut Pasteur, Paris, France; Transcriptome & Epigenome Platform, Genomes & Genetics Department and Center For Innovation and Technological Research, Institut Pasteur, Paris, France

## Abstract

**Background:** Several R packages exist for the detection of differentially expressed genes from RNA-Seq data. The analysis process includes three main steps, namely normalization, dispersion estimation and test for differential expression. Quality control steps along this process are recommended but not mandatory, and failing to check the characteristics of the dataset may lead to spurious results. In addition, normalization methods and statistical models are not exchangeable across the packages without adequate transformations the users are often not aware of. Thus, dedicated analysis pipelines are needed to include systematic quality control steps and prevent errors from misusing the proposed methods.

**Results:** SARTools is an R pipeline for differential analysis of RNA-Seq count data. It can handle designs involving two or more conditions of a single biological factor with or without a blocking factor (such as a batch effect or a sample pairing). It is based on DESeq2 and edgeR and is composed of an R package and two R script templates (for DESeq2 and edgeR respectively). Tuning a small number of parameters and executing one of the R scripts, users have access to the full results of the analysis, including lists of differentially expressed genes and a HTML report that (i) displays diagnostic plots for quality control and model hypotheses checking and (ii) keeps track of the whole analysis process, parameter values and versions of the R packages used.

**Conclusions:** SARTools provides systematic quality controls of the dataset as well as diagnostic plots that help to tune the model parameters. It gives access to the main parameters of DESeq2 and edgeR and prevents untrained users from misusing some functionalities of both packages. By keeping track of all the parameters of the analysis process it fits the requirements of reproducible research.

## Background

DESeq2 [1] and edgeR [2] are very popular Bioconductor [3] packages for differential expression analysis of RNA-Seq, SAGE-Seq, ChIP-Seq or HiC count data. They are very well documented and easy-to-use, even for inexperienced R users. In recent years edgeR and a previous version of DESeq2, DESeq [4], have been included in several benchmark studies [5, 6] and have shown to perform well in replicated experiments. However, running these packages, users can analyse their own dataset without entering important steps of the analysis process such as controlling the quality of the data, exploring its structure or checking some hypotheses of the statistical model. Although these steps are strongly recommended by the authors of the packages [7], they can be skipped by the users without stopping the analysis process.

In an attempt to provide a user friendly access to the whole analysis process of RNA-Seq data, the RNASeqGUI R package has been proposed to provide users with a graphical interface and help R beginners to run a differential analysis without writing R code [8]. The analysis is divided into six main steps among which the pre-analysis section proposes no less than 12 possible figures to explore the data. Similarly, systemPipeR is a R package that proposes a pipeline to process raw fastq RNA-Seq reads from quality filtering to the alignment and counting and to perform a differential analysis using either the edgeR or DESeq2 package [9].

Normalization is another key issue when looking for differentially expressed (DE) genes [10]. The normalization methods associated with edgeR and DESeq2 have been shown to outperform other methods, in particular when expressed RNA repertoires vary across biological conditions or in the presence of highly expressed genes [11]. Although both methods are based on the hypothesis that most genes are not DE, they are computed differently and are not exchangeable unaltered. The size factors computed by DESeq2 are used in the estimation of the mean of the Negative Binomial distribution when the scaling factors proposed by edgeR apply to library sizes (total counts), the normalized library sizes being included as an offset in the statistical model. Thus, an adequate transformation is needed when willing to use any of these normalization methods with any test for differential expression, otherwise leading to inconsistent results as shown in [12] [see Additional file 1].

Similarly, transformation is an important issue at the quality control step where clustering methods and Principal Component Analysis can be used to explore the structure of the dataset. These methods make the assumption that the data are homoscedastic, meaning that the variance is independent of the signal intensity. However count data are not homoscedastic. Using either raw counts or popular transformations such as Reads Per Kilobase per Million mapped reads (RPKM) normalized expression values is not suited for such methods [1, 4].

Finally, in the context of developing reproducible research it is essential to fully describe and explore the data and keep track of the full analysis process with the associated parameter values and software versions. edgeR and DESeq2 do not provide an automatic reporting tool for this purpose.

## Implementation

SARTools (Statistical Analysis of RNA-Seq data Tools) addresses these limitations by proposing a comprehensive, easy-to-use, DESeq2- and edgeR-based R pipeline that covers all the steps of a differential analysis, from the quality control of raw count data to the detection of DE genes. It applies to experimental designs involving one biological factor with two or more levels, such as time series or KO vs. WT experiments. When more than two levels are included in the design, all pairwise comparisons are performed. A blocking factor can be specified to take into account data pairing or the presence of a batch effect (e.g. day of preparation effect). However, SARTools does not handle complex experimental designs with interactions since it involves a careful definition of the design formula and of the contrasts to be tested according to the biological question under study. Indeed, it is neither desired nor safe to automate this part of the analysis process. Users who would have to analyse complex experimental designs are encouraged to use directly either DESeq2 or edgeR which both provide extensive help about this kind of experiments.

SARTools is composed of an R package and two R script templates that allow to run the analysis with either DESeq2 or edgeR. Both scripts rely on each package-specific functions as often as possible, and on SARTools functions to export figures and tables and to generate the HTML report. Each script starts with a section of about 15 parameters that refer to (i) paths to input files and the working directory where the analysis will be performed, (ii) project identification, (iii) experimental design, (iv) normalization and statistical test, (v) filtering process and (vi) plotting. Parameters (i) to (iii) have to be adapted to each analysis. The other parameters have default values and can be left unchanged but are accessible to advanced users if they wish to tune the analysis or the reporting more finely.

SARTools requires two types of input files: count data files containing raw counts and a target file that describes the experimental design [13]. Count data files are sample-specific and are composed of two columns (a unique feature identifier and a raw feature count) with no header. Note that the alignment and counting steps are out of the scope of SARTools and have to be carried out before using specific tools. HTSeq-count output files can be used as input for instance [14]. The target file contains one row per sample and at least three columns with headers: a unique sample identifier or label, the name of the associated raw counts file and the sample biological condition (see Table 1). If a blocking factor has to be accounted for (e.g. in case of batch effect or paired samples), it is reported in a fourth column. These input files are read by SARTools to build a matrix of integer values in which the intersection of the *i*-th row and the *j*-th column reflects how many reads have been mapped to feature *i* in sample *j*. This matrix is then used as input for DESeq2 or edgeR.

**Table 1:**
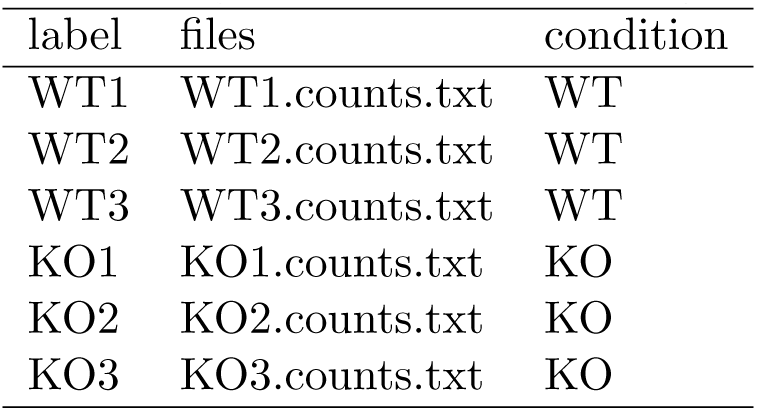
Example of target file for a KO vs. WT experiment including 3 replicates in each biological condition. This table should go at the end of the implementation section.

The source code of the package is available on GitHub (see Availability and requirements section) and is provided in the Additional file 2 as well. Figure 1 describes the successive steps of the workflow and provides the names of the scripts and R functions corresponding to each step. Furthermore, the instructions to integrate SARTools into a Galaxy instance [15, 16, 17] are available on the France Bioinformatique Tool Shed^1^. Galaxy is known to be very user-friendly for biologists and allows them to create worflows to deal with RNA-Seq data. Many tools were already available for the cleaning, mapping and counting steps and SARTools now offers the possibility to run the differential analysis step within the Galaxy environment.

**Figure 1:**
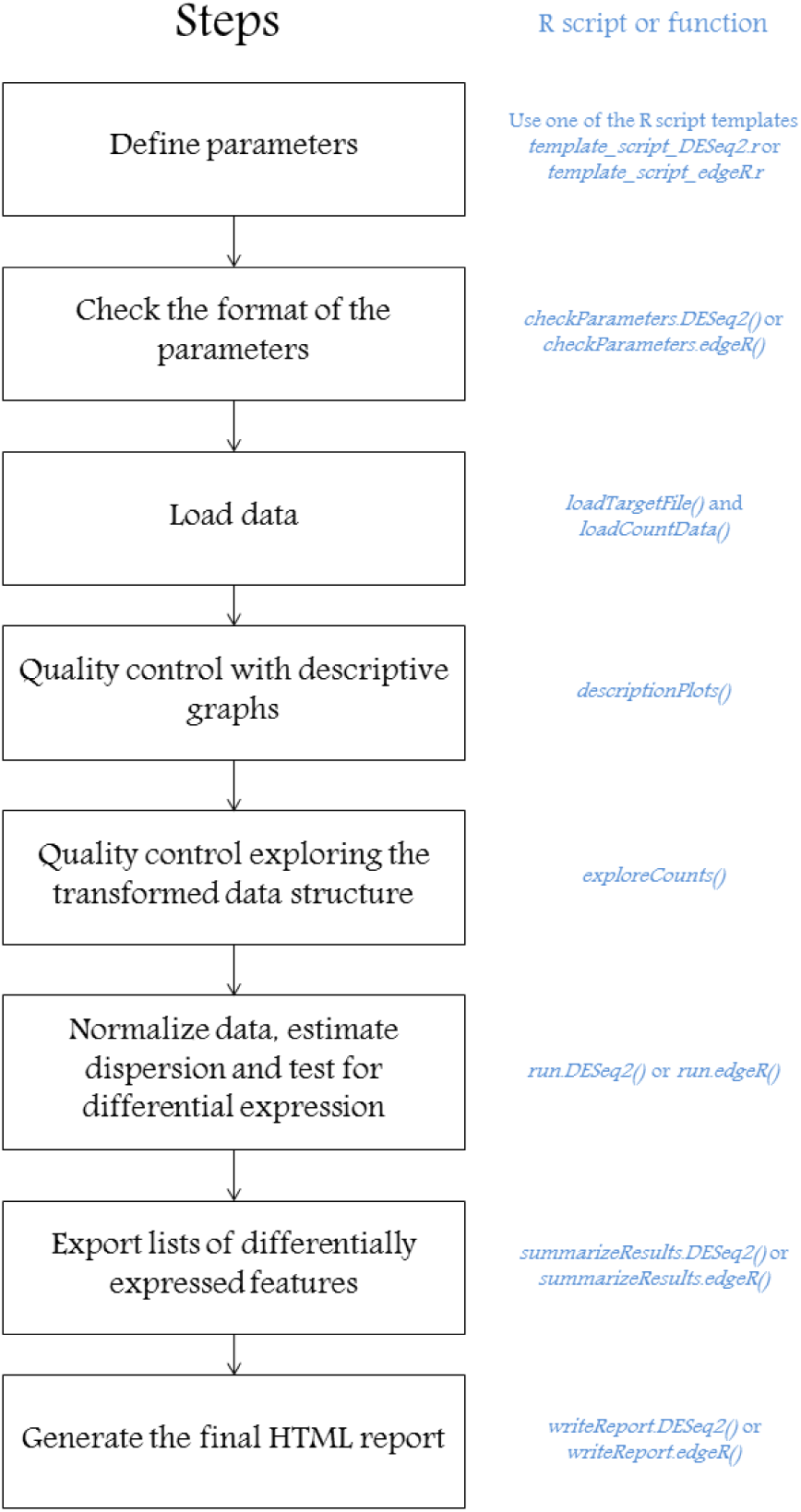
SARTools workflow: Left part (black): successive steps of the analysis; Right part (blue): SARTools functions associated with each step when running either DESeq2 or edgeR.

## Results and Discussion

As soon as the script parameters have been given proper values, and assuming that input files fulfill the format requirements described above, the R script can be run. Using a standard desktop computer, the running time is usually lower than one minute to perform the analysis of 10 samples containing about 50 000 features each. Two types of output files are then made available:

1. a HTML report that describes the whole analysis process in four main steps:

- *data quality control:* numerous plots are proposed to provide a comprehensive description of the dataset, including, for each sample: total number of reads, proportion of null counts, name of the feature having the highest count and proportion of reads associated with it (see Additional file 3 for a full list of the descriptive plots). Principal Component Analysis (PCA) and sample clustering use properly transformed data (with either the Variance Stabilizing Transformation (VST) or the rlog transformation for DESeq2, or log Count Per Million (CPM) for edgeR). These figures provide redundant yet complementary information that help the user to understand the structure of the data and detect possible problems. Legends and titles are added to all figures. The color associated with each sample remains consistent across all figures to ease the interpretation.
- *normalization:* this section uses the default normalization associated with each package, as they have been shown to perform equally well [11]. Two alternative normalization methods are available with edgeR and are accessible through a normalization parameter. When running DESeq2 specific histograms are plotted to check the relevance of the normalization option (refer to the package vignette for additional information). Note that SARTools does not allow the user to perform other normalizations than the ones proposed with the package used for the analysis. For example, it is impossible to provide DESeq2 or edgeR with a matrix of (rounded) RPKM counts as input or any matrix of already normalized values, as both statistical models only accept raw counts as input.
- *test for differential expression:* results include plots to assess the validity of the statistical test as well as lists of up- and down-regulated features. These figures include a histogram a raw p-values, a MA-plot and a volcano-plot for each pairwise comparison. A detailed description of the tab-delimited text files containing these lists is also provided.
- *R session information and parameter values:* this section keeps track of all the information about the packages, methods and parameter values used to run the analysis. In a reproducible research framework, this section is particularly important if the user has to re-perform the analysis later within the same environment and with the same parameters. Furthermore, providing these pieces of information is also often required for publication purposes.
2. three tab-delimited text files per comparison, containing (i) information about all the genes included in the analysis, (ii-iii) lists of significantly up- and down-regulated features ordered by adjusted p-value. Even if Gene Ontology (GO) term analysis is out of the scope of SARTools, these files can be used as a starting point for such an analysis.

All the figures produced by SARTools are integrated into the HTML report and enable the user to perform quality control of the dataset in several ways. Indeed, as they provide complementary pieces of information, they allow a reliable detection and interpretation of possible problems within the dataset and the analysis results. Common problems that may arise from RNA-Seq experiments can be, but are not limited to, the presence of outlier samples, the presence of a batch effect, an inversion between two samples, the presence of a highly expressed sequence in some or all samples, etc. As an example, the presence of an outlier sample in the dataset may be detected in several ways (Figure 2): it can have a much lower - or higher - total number of reads than the other samples. This total count difference may be associated with a higher proportion of null counts (in that case the DESeq2 normalization process will hardly compensate for the difference for these genes having a null count in this sample only, since it is a multiplicative correction), or a clear difference in the density distribution of the counts, or a higher proportion of reads associated with the most represented feature. This outlier sample will also be detected on the PCA plot as well as the clustering one, and may lead to inconsistent values of the SERE coefficients [18]. These complementary information will help to decide whether this sample should be excluded from further analyses or not. The package vignette provides other detailed examples of possible problems that may be encountered and the different plots that can help to interpret inconsistencies between samples (see Additional file 3).

**Figure 2:**
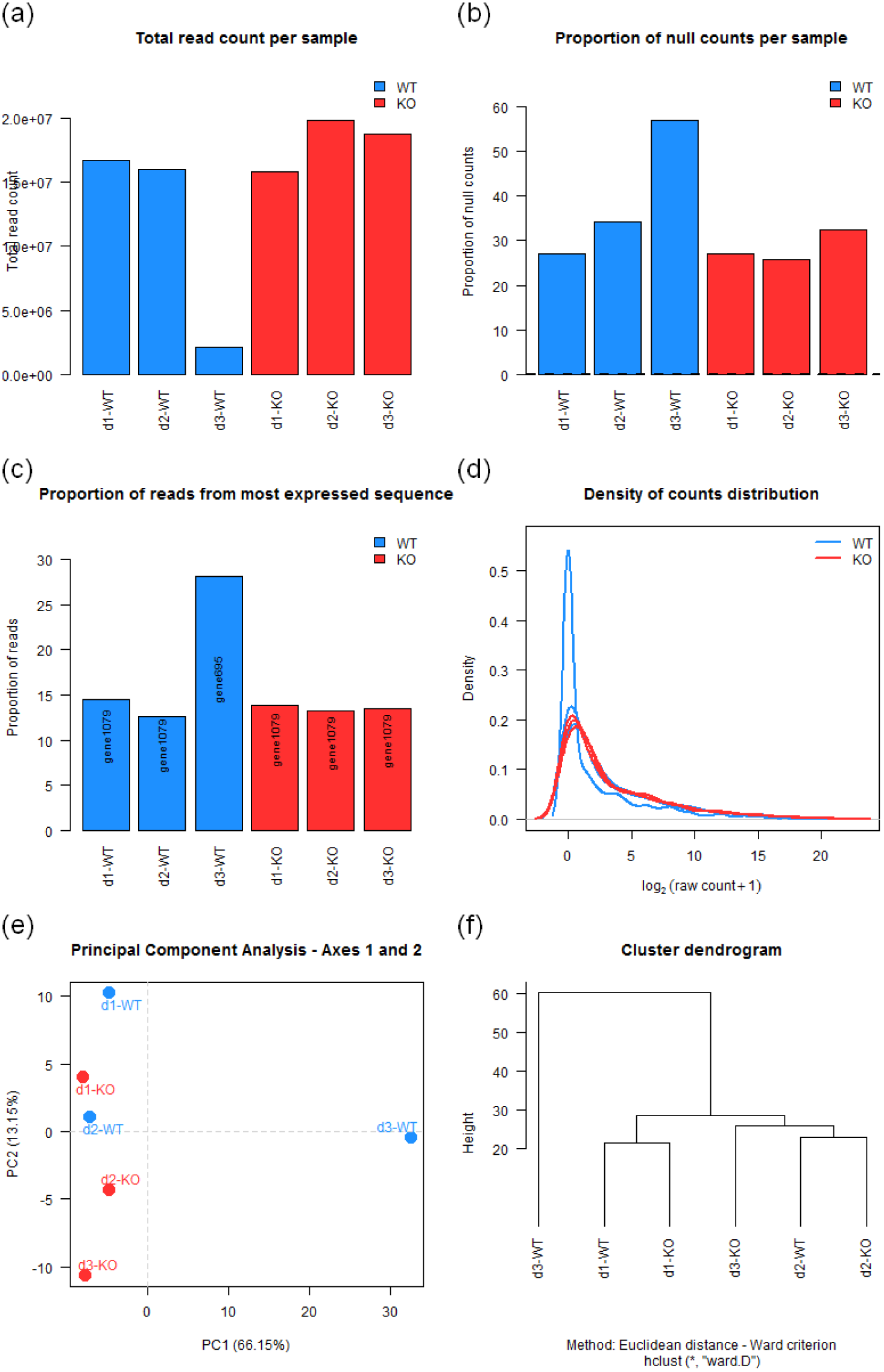
Several ways to detect an outlier (d3-WT). (a) Low total number of reads for d3-WT, (b) High proportion of null counts for d3-WT, (c) Most expressed sequence different for d3-WT, (d) Different distribution of counts for d3-WT, (e-f) d3-WT falls far from the other samples on the first component of the PCA and on the dendrogram.

## Conclusions

SARTools supplies a detailed description of the dataset and proposes diagnostic plots for each step of the analysis. The simple structure of the scripts allow the user to easily and quickly adapt the analysis process to the characteristics of the dataset and to detect possible problems. Unlike other RNA-Seq data analysis pipelines, it does not allow the user to pipe any possible method at any step of the analysis process to build his own workflow, as it focuses on the proper use of these statistical methods. Experienced data analysts may be able to use appropriate command lines to check the quality of raw reads and map them appropriately, they may be regular R users as well without being aware of specific requirements of the statistical models involved and possible problems arising from mixing different methods relying on different models or assumptions. In this context, SARTools provides a safe environment for differential expression analysis of RNA-Seq data on a gene basis.

## Availability and requirements

**Project name:** SARTools.

**Project home page:** https://github.com/PF2-pasteur-fr/SARTools/.

**Operating systems:** Platform independent.

**Programming language:** R.

**Other requirements:** DESeq2^b^, edgeR^c^, knitr^d^, genefilter^e^ and xtable^f^ R packages. All the instructions to install them are given in the README file available on the SARTools GitHub home page.

**License:** GNU General Public License, version 2 (GPL-2.0).

## Competing interests

The authors declare that they have no competing interests.

## Author’s contributions

HV and MAD created the SARTools R package. HV, JYC and MAD wrote this manuscript. All authors read and approved the final manuscript.

## Acknowledgements

We thank Odile Sismeiro, Caroline Proux, Bernd Jagla, Andrea Rau and Christelle Hennequet-Antier for their support and very helpful comments. This work was supported by the INFRASTRUCTURES NATIONALES DE BIOLOGIE ET SANTE France Gnomique (ANR Grant 10- INBS-09-09).

## Additional Files

**Additional file 1 — why are normalization methods not interchangeable?**

This additional file shows why normalization methods are not interchangeable between statistical models without adequate transformation.

**Additional file 2 — Source code of SARTools**

This additional file is an archive (.tar.gz) of the source code of the current version of SARTools (version 1.1.0).

**Additional file 3 — Vignette of the package (version 1.1.0)**

This additional file is the vignette (tutorial.html) distributed with the SARTools package. It provides information on how to use the package and the potential problems that can occur during an analysis. It also shows how to run a toy example to test the pipeline.

## Footnotes

1 http://toolshed.france-bioinformatique.fr/repository?repository_id=529fd61ab1c6cc36&changeset_revision=b73818435a7f

## References

[1] Love M, Huber W, Anders S. Moderated estimation of fold change and dispersion for RNA-Seq data with DESeq2. Genome Biology. 2014; doi:10.1186/s13059-014-0550-8.

[2] Robinson M, McCarthy DJ, Smyth GK. edgeR: a Bioconductor package for differential expression analysis of digital gene expression data. Bioinformatics. 2009; doi:10.1093/bioinformatics/btp616.

[3] Gentleman RC, Carey VJ, Bates DM, et al. Bioconductor: Open software development for computational biology and bioinformatics. Genome Biology. 2004; doi:10.1186/gb-2004-5-10-r80.

[4] Anders S, Huber W. Differential expression analysis for sequence count data. Genome Biology. 2010; doi:10.1186/gb-2010-11-10-r106.

[5] Soneson C, Delorenzi M. A comparison of methods for differential expression analysis of RNA-seq data. BMC Bioinformatics. 2013; doi:10.1186/1471-2105-14-91.

[6] Rapaport F, Khanin R, Lyang Y, et al. Comprehensive evaluation of differential gene expression analysis methods for RNA-seq data. Genome Biology. 2013; doi:10.1186/gb-2013-14-9-r95.

[7] Anders S, McCarthy DJ, Chen Y, et al. Count-based differential expression analysis of RNA sequencing data using R and Bioconductor. Nature Protocols. 2013; doi:10.1038/nprot.2013.099.

[8] Russo F, Angelini C. RNASeqGUI: a GUI for analysing RNA-Seq data. Bioinformatics. 2014; doi:10.1093/bioinformatics/btu308

[9] Girke T. systemPipeR: NGS workflow and report generation environment. R package version 1.0.0. 2014; https://github.com/tgirke/systemPipeR.

[10] Bullard JH, Purdom E, Hansen KD, et al. Evaluation of statistical methods for normalization and differential expression in mRNA-Seq experiments. BMC Bioinformatics. 2010; doi:10.1186/1471-2105-11-94.v

[11] Dillies MA, Rau A, Aubert J, et al. A comprehensive evaluation of normalization methods for Illumina high-throughput RNA sequencing data analysis. Briefings in Bioinformatics. 2013; doi:10.1093/bib/bbs046.

[12] Zhou X, Oshlack A and Robinson MD. miRNA-seq normalization comparisons need improvement. RNA. 2013; doi:10.1261/rna.037895.112.

[13] Ritchie ME, Phipson B, Wu D, et al. limma powers differential expression analyses for RNA-sequencing and microarray studies. Nucleic Acids Research. 2015; doi:10.1093/nar/gkv007.

[14] Anders S, Pyl TP, Huber W. HTSeq - A Python framework to work with high-throughput sequencing data. Bioinformatics. 2014; doi:10.1093/bioinformatics/btu638.

[15] Goecks J, Nekrutenko A, Taylor J and The Galaxy Team. Galaxy: a comprehensive approach for supporting accessible, reproducible, and transparent computational research in the life sciences. Genome Biology. 2010; doi:10.1186/gb-2010-11-8-r86

[16] Blankenberg D, Von Kuster G, Coraor N, et al. Galaxy: a web-based genome analysis tool for experimentalists. Current Protocols in Molecular Biology. 2010; doi:10.1002/0471142727.mb1910s89

[17] Giardine B, Riemer C, Hardison RC, et al. Galaxy: a platform for interactive large-scale genome analysis. Genome Research. 2005; doi:10.1101/gr.4086505

[18] Schulze SK, Kanwar R, Gölzenleuchter M, et al. SERE: Single-parameter quality control and sample comparison for RNA-Seq. BMC Genomics. 2012; doi:10.1186/1471-2164-13-524.

